# GeneTEFlow: A Nextflow-based pipeline for analysing gene and transposable elements expression from RNA-Seq data

**DOI:** 10.1101/2020.04.28.065862

**Authors:** Xiaochuan Liu, Jadwiga R Bienkowska, Wenyan Zhong

**Affiliations:** Oncology Research & Development, Pfizer Worldwide Research and Development, Pearl River, NY 10965, USA; Oncology Research & Development, Pfizer Worldwide Research and Development, San Diego, CA 92121, USA

## Abstract

Transposable elements (TEs) are mobile genetic elements in eukaryotic genomes. Recent research highlights the important role of TEs in the embryogenesis, neurodevelopment, and immune functions. However, there is a lack of a one-stop and easy to use computational pipeline for expression analysis of both genes and locus-specific TEs from RNA-Seq data. Here, we present GeneTEFlow, a fully automated, reproducible and platform-independent workflow, for the comprehensive analysis of gene and locus-specific TEs expression from RNA-Seq data employing Nextflow and Docker technologies. This application will help researchers more easily perform integrated analysis of both gene and TEs expression, leading to a better understanding of roles of gene and TEs regulation in human diseases. GeneTEFlow is freely available at https://github.com/zhongw2/GeneTEFlow.

## Introduction

Transposable elements (TEs) are mobile DNA sequences which have the capacity to move from one location to another on the genome[1]. TEs make up a considerable fraction of most eukaryotic genomes and can be classified into retrotransposons and DNA transposons according to their different mechanisms of transposition and chromosomal integration[2, 3]. Retrotransposons are made of Long Terminal Repeats (LTRs) and non-LTRs that include long interspersed nuclear elements (LINEs) and short interspersed nuclear elements (SINEs) that mobilize via a RNA intermediate, while DNA transposons mobilize and function through a DNA intermediate[4–6]. TEs can be transcribed from the genome[7] and have been demonstrated to play important roles in the mammalian embryogenesis[8, 9], neurodevelopment[10, 11], and immune functions[12, 13]. Furthermore, aberrant expressions of TEs have been linked to cancers[14–16], neurodegenerative disorders[17, 18], and immune-mediated inflammation[19, 20]. Therefore, it has become increasingly important to explore biological roles of TEs expression. However, genome-wide analysis of TEs expression from high throughput RNA sequencing data has been a challenging computational problem. TEs contain highly repetitive sequence elements, making it arduous to unambiguously assign reads to the correct genomic location and accurately quantitate their expression level. Several bioinformatics tools have been developed to address this challenge with relatively good success [16, 21–23]. Recently, SQuIRE was reported to have the capability to quantify locus-specific expression of TEs from RNA-Seq data[23]. In addition, RNA-Seq data has long been used to detect dysregulated genes between different disease and/or drug treatment conditions to help understand disease mechanisms and/or drug response mechanisms. Therefore, it is of great interest to quantify both TEs and gene expression to elucidate contribution of both to disease mechanisms. Although many open source software and tools exist for analysing gene [24–26] and TEs expression, there are considerable challenges to efficiently apply these tools. In general, these multi-step data processing pipelines use many different tools. Correct versions of each tool need to be installed separately, and appropriate options, parameters, different reference genome and gene annotation files have to be set at each step. This can be quite tedious and challenging especially for non-computational users. Additionally, to ensure reproducibility of the analysis results, it is critical to capture analysis parameters from each step of the process. Equally important, to enable general use of the pipeline, the pipeline should be platform agnostic. Thus far, a one-stop computational framework for the comprehensive analysis of gene and locus-specific TEs expression from RNA-Seq data does not exist.

To address this need, we developed GeneTEFlow, a reproducible and platform-independent workflow, for the comprehensive analysis of gene and locus-specific TEs expression from RNA-Seq data using Nextflow[27] and Docker[28] technologies. GeneTEFlow provides several features and advantages for integrated gene and TEs transcriptomic analysis. First, by employing Docker technology, GeneTEFlow encapsulates bioinformatics tools and applications of specific versions into Docker containers enabling tracking, eliminating the need for software installation by users, and ensuring portability of the pipeline on multiple computing platforms including stand-alone workstations, high-performance computing (HPC) clusters, and cloud computing systems. Second, GeneTEflow uses Nextflow to define the computational workflows, not only enabling parallelization and complete automation of the analysis, but also providing capability to track analysis parameters. Thus, GeneTEFlow allows users to generate reproducible analysis results through utilization of both Docker and Nextflow in a platform independent manner. Lastly, GeneTEFlow has modular architecture, and modules in GeneTEFlow can be turned on or off, providing developers with flexibility to build extensions tailored to specific analysis needs.

## Implementation

The GeneTEFlow pipeline was developed using Nextflow, a portable, flexible, and reproducible workflow management system, and Docker technology, a solution to securely build and run applications on multiple platforms. To build the GeneTEFlow pipeline, a series of bioinformatics tools (S1 Table) were selected for QC, quantitation and differential expression analysis of genes and TEs from RNA-Seq data. These bioinformatics tools and custom scripts were built into four Docker containers to ensure portability of the workflow on different computational platforms. Data processing and analysis steps were implemented by modules using Nextflow. Modules are connected through channels and can be run in parallel. Each module in GeneTEFlow can include any executable Linux scripts such as Perl, R, or Python. Parameters for each module are defined in a configuration file.

A conceptual workflow of GeneTEFlow is illustrated in Fig 1. The workflow includes four major inputs: raw sequence files in fastq format, a sample meta data file in excel format, reference genome and gene annotation files, and a Nextflow configuration file. The sample meta data file contains detailed sample information and the design of group comparisons between different experimental conditions. Human reference genome UCSC hg38 with the gene annotation (.gtf) was downloaded from Illumina iGenomes collections[29] and used by the bioinformatics tools included in GeneTEFlow. Scheduling of computational resources for each application module is defined in the configuration file.

**Fig 1.**
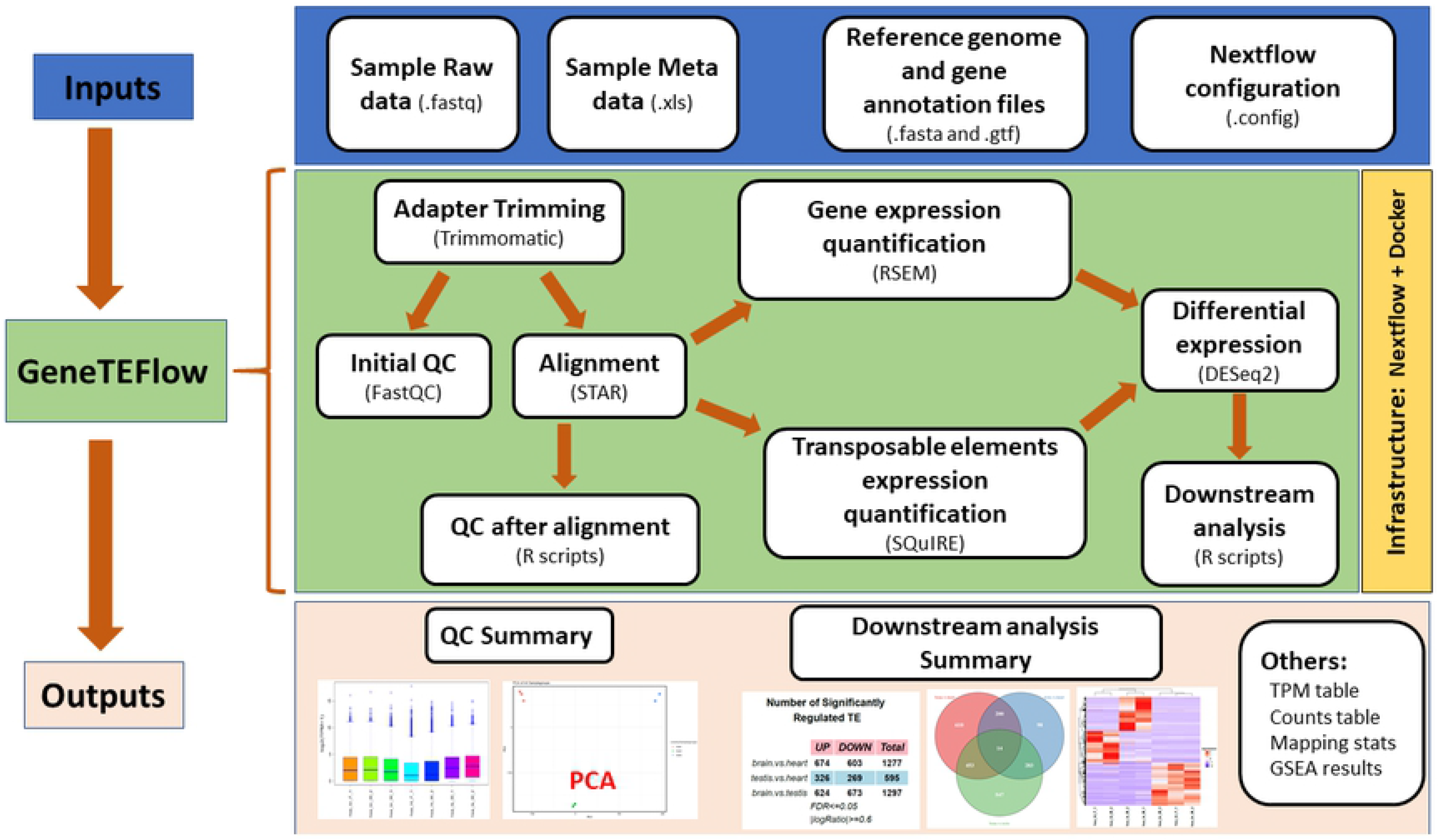
Illustration of GeneTEFlow: a Nexflow-based pipeline for identification of differentially expressed genes and locus specific transposable elements from RNA-Seq data.

GeneTEFlow analysis is performed in following steps: QC, expression quantification, differential expression and down-stream analysis. First, adapter sequences are trimmed off from the Illumina raw reads using Trimmomatic(v0.36)[30] for single-end or paired-end reads, and low-quality reads are filtered out. Next, FastQC(v0.11.7)[31] is executed to survey the quality of sequencing reads, and report is generated to help identify any potential issues of the high throughput sequencing data. Reference genome index for mapping sequencing reads to mRNA genes is built using “rsem-prepare-reference” of RSEM (v.1.3.0). Reads remaining after the pre-processing step are mapped to the reference genome using STAR(v2.6.0c)[32]. Gene level expression is quantitated as expected counts and transcripts per million (TPM) using “rsem-calculate-expression” of RSEM(v1.3.0) with default parameters [33]. Custom Perl scripts were developed to aggregate data from each sample into a single data matrix for expected counts and TPM values respectively. The expression quantification of locus-specific TEs is performed by SQuIRE[23].

In addition, we also implemented quality control measures after reads alignment step to detect potential outlier samples resulted from experimental errors. Boxplot and density plot are used to evaluate the overall consistency of the expression distribution for each sample. Sample correlation analysis is performed with Pearson method using TPM values to assess the correlation between biological replicates from each sample group. Principal component analysis (PCA) is employed to identify potential outlier samples and to investigate relationships among sample groups.

Differential expression analysis of genes and transposable elements is performed using DESeq2(v1.18.1) package[34]. Significantly up-regulated and down-regulated genes and TEs are summarized in a table. To analyse overlap among significantly regulated genes and TEs from pair-wise comparisons between different sample groups we use Venn diagrams. We perform hierarchical clustering of significantly dysregulated genes or TEs using R package “ComplexHeatmap” [35] with euclidean distance and average linkage clustering parameters. Gene set enrichment analysis (GSEA, v3.0) [36] is conducted using collections from the Molecular Signatures Database (MSigDB) [37]. The outputs (S2 Table) from GeneTEFlow are organized into several folders predefined in a GeneTEFlow configuration file. A tutorial with detailed instructions on how to set up and run GeneTEFlow is provided at https://github.com/zhongw2/GeneTEFlow

## Application of GeneTEFlow

We applied GeneTEFlow to a public dataset from Brawan’s study [38] investigating tissue-specific expression changes of genes and transposable elements. Human RNA-Seq data from brain, heart and testis tissues were downloaded from GEO (accession number: GSE30352) (S3 Table). Expression analysis of genes and TEs were performed using GeneTEFlow and results are shown in Fig 2. Gene expression analysis was performed using RSEM and DESeq2 modules while TEs expression analysis was conducted using SQuIRE and DESeq2 modules within GeneTEFlow. Significantly regulated genes were identified with FDR less than 0.05 and fold change greater than 2. Significantly regulated locus-specific transposable elements were identified with FDR less than 0.05 and fold change greater than 1.5. The number of significantly regulated genes and transposable elements were summarized into two tables respectively (Fig 2, top panels). Using GeneTEFlow, we detected genes and TE differentially expressed between different tissue types (brain vs heart tissues: 6,264 genes and 1,277 TEs; testis vs heart tissues: 7,066 genes and 595 TE; brain vs testis tissues: 8,125 genes and 1,297 TEs) with most significant gene and TE expression differences observed being between brain and testis tissues. Our analysis identified large number of both genes and TEs with tissue specific patterns (Fig 2, middle panels and bottom panels). More in depth analysis to include additional tissue types would be required to fully understand the tissue specific gene and TEs expression and their relationship. GeneTEFlow is a computational solution to facilitate such studies.

**Fig 2.**
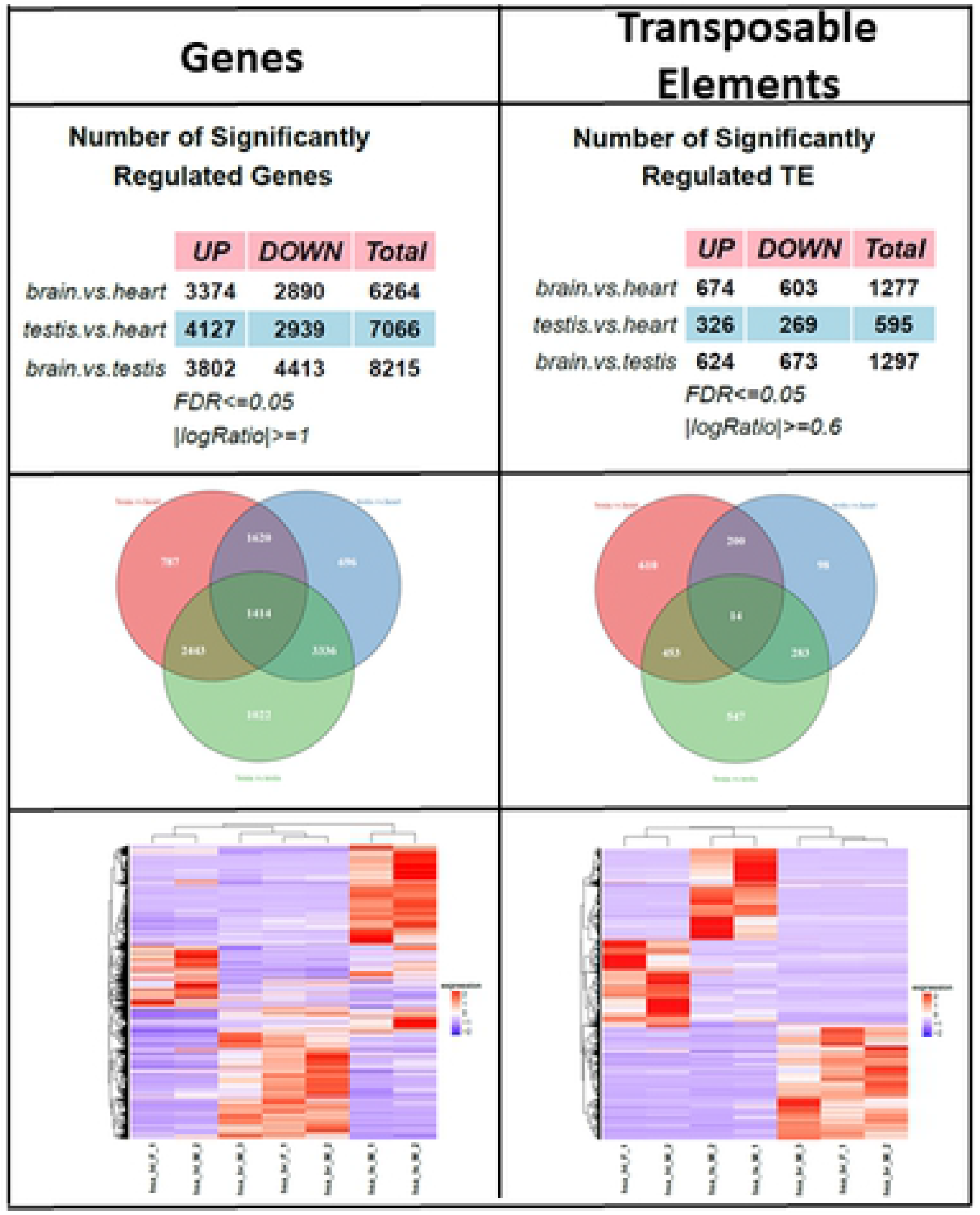
Differential expression analysis results of genes and transposable elements from GeneTEFlow. Left panels: gene results; right panels: TEs results. Top panels: number of significantly regulated genes or TEs in each sample group comparison. Significance was defined as following: FDR ≤ 0.05 and fold change ≥ 2 for gene expression analysis; FDR ≤ 0.05 and fold change ≥ 1.5 for TEs expression analysis. Middle panels: overlaps of significantly regulated genes or TEs amongst sample group comparisons. Bottom panels: hierarchical clustering of significantly regulated genes or TEs.

In addition to quantification of TEs expression, SQuIRE provides quantification of gene expression. Therefore, we compared gene level expression quantification between RSEM and SQuIRE (S1 Fig). The results showed high concordance (correlation coefficient: ~97%) of the gene level expression quantification between the two methods (S1 Fig, highlighted in red box) suggesting a robust measurement for both gene and TEs expression by SQuIRE.

## Conclusions

In conclusion, we have developed and made available an automated pipeline to comprehensively analyse both gene and locus-specific TEs expression from RNA-Seq data. Taking advantage of the advanced functionalities provided by Nextflow and Docker, GeneTEFlow allows users to run analysis reproducibly on different computing platforms without the need for individual tool installation and manual version tracking. We believe this pipeline will be of great help to further our understanding of roles of both gene and TEs regulation in human diseases. This pipeline is flexible and can be easily extended to include additional types of analysis such as alternative splicing, fusion genes, and so on.

## Competing interests

WZ and JRB are employees of Pfizer Inc.

XL was contractor of Pfizer Inc. when the work was being conducted.

## Authors’ contributions

WZ conceptualized the work. XL and WZ designed and implemented the pipeline. XL, JRB and WZ drafted and revised the manuscript. All authors read and approved the final manuscript.

## Acknowledgements

We gratefully acknowledge inputs and support from our colleagues: Jeremy Myers, Keith Ching, Corey Dasilva and Da Tse.

## Supporting information

**S1 Fig**. Comparison of gene expression quantification by RSEM and SQuIRE. Gene expression (total 22,955 genes) of samples from brain tissues (left), heart tissues (middle), and testis tissues (right) was calculated by both RSEM and SQuIRE. Lower diagonal panels: pairwise comparisons using log2(TPM + 1) of 22,955 genes. Upper diagonal panels: correlation coefficient of each comparison. Panels highlighted in red: correlation coefficient of comparisons between RSEM and SQuIRE gene expression quantification of the same sample. Rep_: replicate, _RSEM: quantification performed by RSEM, _SQuIRE: quantification performed by SQuIRE.

**S1 Table**. Major bioinformatics tools installed in GeneTEFlow

**S2 Table**. Major outputs from GeneTEFlow

**S3 Table**. Human RNA-Seq data used in the example application of GeneTEFlow

**S1_File**. Supplemental tables: S1-S3 Tables.

